# A framework to efficiently smooth *L*_1_ penalties for linear regression

**DOI:** 10.1101/2020.09.17.301788

**Authors:** Georg Hahn, Sharon M. Lutz, Nilanjana Laha, Christoph Lange

## Abstract

Penalized linear regression approaches that include an *L*_1_ term have become an important tool in statistical data analysis. One prominent example is the least absolute shrinkage and selection operator (Lasso), though the class of *L*_1_ penalized regression operators also includes the fused and graphical Lasso, the elastic net, etc. Although the *L*_1_ penalty makes their objective function convex, it is not differentiable everywhere, motivating the development of proximal gradient algorithms such as Fista, the current gold standard in the literature. In this work, we take a different approach based on smoothing in a fixed parameter setting (the problem size *n* and number of parameters *p* are fixed). The methodological contribution of our article is threefold: (1) We introduce a unified framework to compute closed-form smooth surrogates of a whole class of *L*_1_ penalized regression problems using Nesterov smoothing. The surrogates preserve the convexity of the original (unsmoothed) objective functions, are uniformly close to them, and have closed-form derivatives everywhere for efficient minimization via gradient descent; (2) We prove that the estimates obtained with the smooth surrogates can be made arbitrarily close to the ones of the original (unsmoothed) objective functions, and provide explicitly computable a priori error bounds on the accuracy of our estimates; (3) We propose an iterative algorithm to progressively smooth the *L*_1_ penalty which increases accuracy and is virtually free of tuning parameters. The proposed methodology is applicable to a large class of *L*_1_ penalized regression operators, including all the operators mentioned above. Although the resulting estimates are typically dense, sparseness can be enforced again via thresholding. Using simulation studies, we compare our framework to current gold standards such as Fista, glmnet, gLasso, etc. Our results suggest that our proposed smoothing framework provides predictions of equal or higher accuracy than the gold standards while keeping the aforementioned theoretical guarantees and having roughly the same asymptotic runtime scaling.

## 1 Introduction

In this communication, we aim to efficiently solve *L*_1_ penalized regression problems. Since first considered around 1800 by Adrien-Marie Legendre and Carl Friedrich Gauss, the least squares approach has traditionally been used to solve linear regression problems. Nevertheless, least squares estimates have two major drawbacks, a lack of robustness and a lack of sparsity in high dimensional settings, i.e. settings characterized by *p* ≫ *n*, where *p* ∈ ℕ denotes the number of covariates and *n* ∈ ℕ is the sample size. Those two limitations have motivated the development of new estimation operators that are more suitable for such settings, e.g. the *least absolute shrinkage and selection operator* (Lasso) (Tibshirani, 1996), or the *least-angle regression* (LARS) (Efron et al., 2004). Many important extensions of the standard Lasso approach have been proposed, such as the *elastic net* (Zou & Hastie, 2005), *the fused Lasso* (Tibshirani et al., 2005), *or the graphical Lasso* (Friedman et al., 2008). All Lasso approaches involve an *L*_1_ penalty on the parameters which are being estimated to enforce sparseness.

This article focuses on penalized *L*_1_ regression, where we model the relationship of a given design matrix *X* ∈ ℝ^*n*×*p*^ to an observed response *y* ∈ ℝ^*n*^ with the help of some unknown parameters *β* ∈ ℝ^*p*^. In the most general case, we consider any regression operator which can be written as

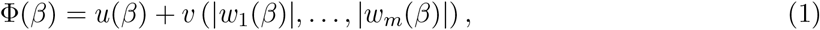

where *u* : ℝ^*p*^ → ℝ implicitly depends on *X* and *y, v* : ℝ^*m*^ → ℝ, and *w*_*j*_ : ℝ^*p*^ → ℝ for *m* ∈ ℕ and *j* ∈ {1, *…, m*}. The regression estimate is computed accordingly as arg min_*β*_ Φ(*β*). Throughout the article, we consider a fixed parameter setting (*n, m, p* are fixed).

Amongst others, all the previously mentioned estimation operators can be casted in the form of eq. (1). For instance, for a fixed design matrix *X* ∈ ℝ^*n*×*p*^ and response *y* ∈ ℝ^*n*^, the objective function of the standard Lasso (Tibshirani, 1996) is obtained from eq. (1) by choosing 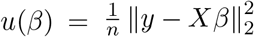, as well as 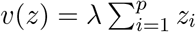 for *z* ∈ ℝ^*p*^, and *w*_*j*_(*β*) = *β*_*j*_ for *j* ∈ {1, *…, p*}.

If the objective function in eq. (1) is convex, steepest descent (quasi-Newton) methods can be employed to compute arg min_*β*_ Φ(*β*). Nevertheless, the non-differentiability of the *L*_1_ penalty may cause a loss of accuracy in conventional gradient(-free) solvers.

We address this issue by providing a smooth surrogate Φ^*μ*^ of eq. (1), which depends on a smoothing parameter *μ >* 0. The smooth surrogate is derived by applying Nesterov smoothing (Nesterov, 2005) to the absolute values in eq. (1). Under the condition that *u* is differentiable and convex, *v* is differentiable and Lipschitz continuous, *v* preserves strict convexity, and *w*_*j*_ are differentiable for *j* ∈ {1, *…, p*}, our framework provides the following guarantees:

1. The surrogate Φ^*μ*^ has explicit gradients everywhere.
2. The surrogate Φ^*μ*^ is strictly convex.
3. The surrogate Φ^*μ*^ is uniformly close to Φ, in the sense that it satisfies 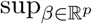 |Φ(*β*) − Φ^*μ*^(*β*)| ≤*C*_*μ*_ = *O*(*μ*) for fixed *m*, where *C*_*μ*_ is a constant which does not depend on *β*.
4. We prove explicit bounds on ∥ arg min_*β*_ Φ(*β*) − arg min_*β*_ Φ^*μ*^(*β*) ∥_2_, that is on the distance between the minimizers of the unsmoothed objective function and its smooth surrogate.
5. We prove that ∥ arg min_*β*_ Φ(*β*) − arg min_*β*_ Φ^*μ*^(*β*) ∥ → 0 in the supremum norm as *μ* → 0.

In particular, all these properties hold true for the aforementioned regression operators, thus making it possible to obtain a unified surrogate with the theoretical guarantees enumerated above for a class of common regression operators. Although the estimates obtained by minimizing Φ^*μ*^ are typically dense, sparseness can be enforced again via thresholding.

To summarize, the contribution of our article is threefold: (1) We introduce the closed-form surrogate Φ^*μ*^; (2) We prove the theoretical properties of Φ^*μ*^ enumerated above; (3) Starting with a high degree of smoothness (i.e., a large value of *μ*), an iterative algorithm is proposed to progressively smooth the surrogate in order to facilitate minimization. The choice of the smoothing parameter does not have a major effect on the performance of our progressive smoothing algorithm, thus making it virtually free of tuning parameters.

We evaluate our proposed algorithms with respect to accuracy and runtime using simulation studies. Our aim is to compare them to the current gold standards in the literature. First, we use as a benchmark the implementation of the Fista algorithm (Beck & Teboulle, 2009) that is included in the *fasta* R-package on CRAN (Chi et al., 2018). Fista combines the basic iterations of the Iterative Shrinkage-Thresholding Algorithm (Daubechies et al., 2004) with a Nesterov acceleration step. The algorithm can be regarded as a proximal gradient version of the Nesterov method (Nesterov, 1983). Analogously to eq. (1), Fista requires the separate specification of the smooth and non-smooth parts of the objective function and their explicit gradients. For the application of our smoothing algorithms to the elastic net, we benchmark against the *glmnet* algorithm (Friedman et al., 2010a) that is implemented in the R-package *glmnet* (Friedman et al., 2020). The glmnet algorithm is a variant of Fista which performs a cyclic update of all coordinates, whereas Fista updates all coordinates per iteration. For the smoothed fused Lasso, we compare with the fused Lasso methodology in the R-package *genlasso* (Arnold & Tibshirani, 2020). Finally, we benchmark our smoothed graphical Lasso against the R-package *glasso* (Friedman et al., 2019).

Since the *least-angle regression* (LARS) (Efron et al., 2004) or the *group Lasso* (Yuan & Lin, 2005) do not involve an *L*_1_ penalty on the regression estimates, our smoothing framework does not apply to them. We provide a detailed literature review in Section 1.1 to highlight previous work, distinguish it from ours, and emphasize the contribution of our article.

This article is structured as follows. Section 2 starts by deriving the smooth surrogate of the objective function. We also derive the aforementioned theoretical guarantees of our smoothing framework, and state precise conditions on the quantities involved in eq. (1) which ensure that the theoretical guarantees hold true. Moreover, we demonstrate how all the aforementioned *L*_1_ penalized regression operators fit in our framework, and we refine our approach by proposing a iterative and virtually tuning-free smoothing procedure. We benchmark our proposed methodology on simulated data against state-of-the-art algorithms in Section 3. The article concludes with a discussion in Section 4. A detailed overview of Nesterov smoothing (Section A), all proofs (Section B), and additional simulations (Section C) are provided in the Supporting Information.

This article extends a previous publication (Hahn et al., 2021) in which Nesterov smoothing is applied to the Lasso objective function only, meaning without any theoretical results. In contrast to the previous work (Hahn et al., 2021), Section 2 presents a framework to smooth an entire class of *L*_1_ penalized regression operators, and the theoretical results of Sections 2.3 and 2.4, the progressive smoothing algorithm of Section 2.5, as well as all simulations of Section 3 are new. The methodology of this article is implemented in the R-package *smoothedLasso* (Hahn et al., 2020), available on *The Comprehensive R Archive Network (CRAN)* (R Core Team, 2014).

Throughout the article, the *i*th entry of a vector *v* is denoted as *v*_*i*_. The entry (*i, j*) of a matrix *X* is denoted as *X*_*ij*_, while the *i*th row of *X* is denoted as *X*_*i*,·_ and the *j*th column is denoted as *X*_·,*j*_. The absolute value, the *L*_1_ norm, as well as the Euclidean norm are written as | · |, ∥ · ∥_1_, and ∥ · ∥_2_, respectively. The Kronecker delta function is written as *δ*_*ij*_, defined as *δ*_*ij*_ = 1 if *i* = *j* and *δ*_*ij*_ = 0 if *i* ≠ *j*. The size of a set *S* is denoted as |*S*|.

### 1.1 Literature review

Since the seminal publication of the Lasso (Tibshirani, 1996), numerous approaches have focused on (smoothing) approaches to facilitate the minimization of the Lasso objective function, or other related objective functions. Nevertheless, the following publications differ from our work in that they do not consider a unified framework to smooth *L*_1_ regularized regression operators, and if they provide a tailored solution to a particular regression operator, the bounds provided differ from the ones we establish on the accuracy of the unsmoothed objective and surrogate. Most importantly, none of the publications we are aware of quantifies the distance between the minimizer of the unsmoothed objective and the minimizer of the surrogate a priori (as we do with our explicitly computable bound), and no progressive smoothing procedure yielding stable regression estimates is derived.

Smoothing approaches for the Lasso *L*_1_ penalty, the SCAD (Smoothly Clipped Absolute Deviation) penalty, and hard thresholding penalties have been considered (Fan & Li, 2001). However, these smoothing approaches are not based on the Nesterov framework (Nesterov, 2005). Instead, the authors employ a quadratic approximation at the singularity of the penalties to achieve a smoothing effect, and they propose a one-step shooting algorithm for minimization. However, their main focus is on root-n consistency results of the resulting estimators and asymptotic normality results for the SCAD penalty, results which the authors state do not all apply to the Lasso.

Some smoothing approaches (Belloni et al., 2011; Chen et al., 2010a; Banerjee et al., 2008) build upon Nesterov’s first-order accelerated gradient descent algorithm (Nesterov, 2005). Those variants of Nesterov’s algorithm are iterative methods which are unrelated to our adaptive smoothing procedure. A detailed overview of several variants of the first-order accelerated gradient descent algorithm can be found in the literature (Becker & Candès, 2011).

Publications which extend the work of Nesterov (Nesterov, 2005) are available (Beck & Teboulle, 2012). In the latter, the authors consider a more general smoothing framework which, as a special case, includes the same smoothing we establish for the absolute value in the *L*_1_ penalty of the Lasso. However, in contrast to our work, the authors do not consider a smooth surrogate of a general form that applies to a wide range of regression operators, and the theoretical guarantees on the surrogate that we derive are unconsidered.

In another publication (Haselimashhadi & Vinciotti, 2016), the authors likewise smooth the absolute value in the *L*_1_ penalty of the Lasso using Nesterov’s technique, and they state the same bound on the difference between the unsmoothed and smoothed Lasso objective functions taken from Nesterov’s results. However, their results focus on the Lasso only, the general surrogate we provide is unconsidered, and no explicit a priori bounds on the accuracy of the obtained minimizers are given. Importantly, these works (Haselimashhadi & Vinciotti, 2016) differ from our work in that the authors enforce that the smoothed Lasso penalty passes through zero, leading the focus of their article to be on another smoothed Lasso approach which is based on the error function of the normal distribution.

Further work available in the literature employs Nesterov’s smoothing techniques for a variety of specialized Lasso objective functions. For instance, some authors (Chen et al., 2010b) consider the group Lasso and employ Nesterov’s formalism to smooth the Lasso penalty using the squared error proximity function which we also consider. Nevertheless, the authors focus on adapting Nesterov’s first-order accelerated gradient descent algorithm in order to compute the Lasso regression estimate, whereas we focus on deriving theoretical bounds on the smooth surrogate and our progressive smoothing algorithm. In our work, we do not propose a modification of Nesterov’s first-order accelerated gradient descent algorithm. The group Lasso is also considered in the literature by some authors (Chen et al., 2012), who separate out the simple nonsmooth *L*_1_ penalty from the more complex structure-inducing penalties, and only smooth the latter. This leaves the *L*_1_ norm on the parameters unchanged, thus still enforcing individual feature level sparsity. In contrast to this published work (Chen et al., 2012), we always smooth all *L*_1_ penalties which are present in the objective function.

The joint Lasso is also considered in the literature (Dondelinger & Mukherjee, 2020), who state an iterative minimization procedure which smoothes the Lasso penalty using Nesterov’s techniques. The authors state closed form derivatives for minimization (as we do as well in the form of the more general derivatives of the smooth surrogate), but no other theoretical results are given.

One variant of the original Lasso which has recently gained attention is the concomitant Lasso. The concomitant Lasso augments the original Lasso with a term *σ/*2 for which a second regularization parameter *σ* is introduced (Ndiaye et al., 2017). The parameter *σ* is meant to be decreased to zero. Smoothing the concomitant Lasso has the advantage that Nesterov’s techniques do not need to be applied to the *L*_1_ penalty. Instead, the smooth concomitant Lasso has a closed form expression which is different from the smoothed Lasso approaches we consider (Ndiaye et al., 2017), and results in the literature (Massias et al., 2018) are only named in analogy to the smoothing terminology introduced in Nesterov’s original paper (Nesterov, 2005). Since the concomitant Lasso does not contain an *L*_1_ penalty, our results to not apply.

## 2 The smooth surrogate

This section states the precise conditions and results of our smoothing framework for regression operators with an objective function of the form of eq. (1). The smoothing of eq. (1) will be achieved with the help of Nesterov smoothing, which is briefly summarized in Section 2.1. In Section 2.2 we smooth the *L*_1_ contribution in eq. (1), leading to the smooth surrogate objective function. Section 2.3 states the theoretical guarantees provided for the smooth surrogate, and the precise conditions required for the guarantees to hold true. Section 2.4 gives examples of how common regression operators fit into our framework. Finally, Section 2.5 refines our approach by proposing a iterative and virtually tuning-free smoothing procedure.

### 2.1 Summary of Nesterov smoothing

The smoothing of a piecewise affine and convex function *f* : ℝ^*q*^ → ℝ, where *q* ∈ ℕ, has been established in the literature (Nesterov, 2005). Since *f* is piecewise affine, it can be represented as *f* (*z*) = max_*i*=1,…,*k*_ (*A*[*z*, 1]^T^) _*i*_, where *k* ∈ ℕ is the number of linear pieces (components). Here, the rows of the matrix *A* ∈ ℝ^*k*×(*q*+1)^ contain the coefficients of each linear piece, with column *q* + 1 containing all constant coefficients. Moreover, [*z*, 1] ∈ ℝ^*q*+1^ denotes the vector obtained by concatenating *z* ∈ ℝ^*q*^ and the scalar 1.

The goal is to derive an approximation *f*^*μ*^ of *f* which is both smooth and uniformly close to *f* (Nesterov, 2005). The approximation *f*^*μ*^ depends on a so-called proximity (or prox) function, see Section A.1, which is parameterized by a smoothing parameter *μ >* 0 controlling the degree of smoothness. We consider two choices of the prox function, presented in Section A.2. The so-called entropy prox function yields a closed-form expression of the smooth approximation 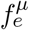 given by

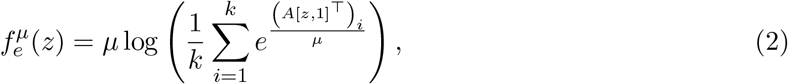

which satisfies the uniform approximation bound

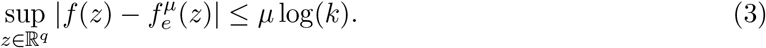

Another choice, introduced in Section A.2, is the squared error prox function, given by 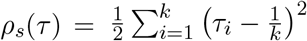 for *τ* ∈ ℝ^*k*^. Let the vector *c*(*z*) = (*c*_1_(*z*), *…, c*_*k*_ (*z*)) ∈ ℝ^*k*^ be defined componentwise by *c*_*i*_(*z*) = 1*/μ* · (*A*[*z*, 1]^T^)_*i*_ − 1*/k* for *i* ∈ {1, *…, k*}. Let *ĉ*(*z*) ∈ ℝ be the Michelot projection (Michelot, 1986) *of c*(*z*) onto the *k*-dimensional unit simplex *Q*_*k*_ (see Section A.2.2). In the original publication (Nesterov, 2005), it is shown that

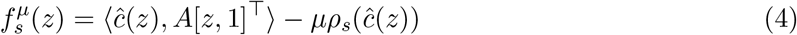

is a smooth approximation of *f* satisfying the uniform bound

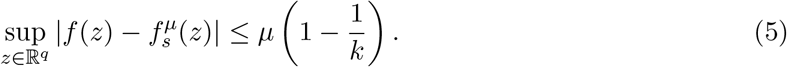

Further details on the above results can be found in Section A of the Supplementary Materials.

### 2.2 The smooth surrogate

We employ the results of Section 2.1 to smooth the absolute value in eq. (1). In the following subsections, we always consider *k* = 2, *z* ∈ ℝ, and the specific choice of the matrix *A* ∈ ℝ^2×2^ given by

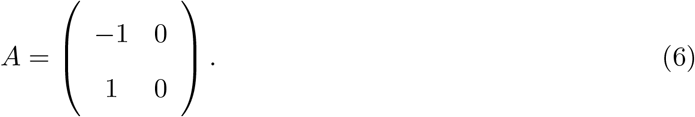

The specific choice of *A* in eq. (6) allows us to express the (one dimensional) absolute value as a piecewise affine and convex function *f* (*z*) = max{−*z, z*} = max_*i*=1,2_ (*A*[*z*, 1]^T^)_*i*_.

Simplifying eq. (2) with the specific choice of *A* of eq. (6) results in the entropy prox approximation of the absolute value and its explicit derivative, given by

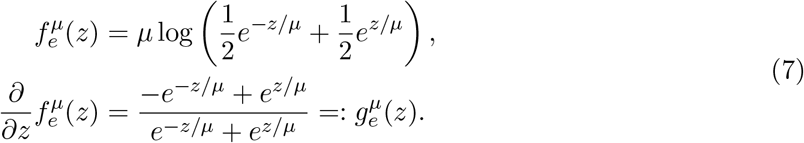

The approximation bound of eq. (3) simplifies to

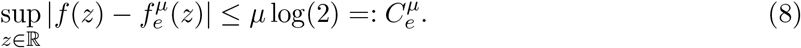

In a similar fashion, the squared error prox smoothing of eq. (4) with *A* as in eq. (6) leads to the following approximation of the absolute value and its explicit derivative:

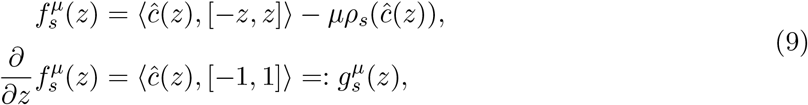

where *ĉ*(*z*) ∈ ℝ^2^ denotes the Michelot projection of *c*(*z*) = 1*/μ* · [−*z, z*] − 1*/*2 onto the twodimensional unit simplex. A derivation of the derivative of 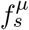 can be found in Lemma 4 (Hahn et al., 2017). The approximation bound of eq. (5) simplifies to

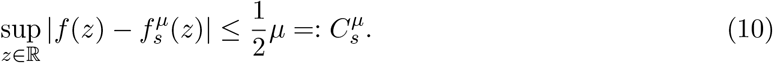

With the smooth approximation of the absolute value in place, we define the surrogate of Φ as

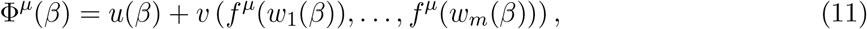

where in the remainder of Section 2, *f*^*μ*^ always denotes either the entropy prox smoothed absolute value 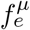, or the squared error prox smoothed absolute value 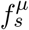.

### 2.3 Theoretical guarantees

Depending on the conditions imposed upon *u, v*, and *w*_*j*_ for *j* ∈ {1, *…, m*} in eq. (1), the surrogate Φ^*μ*^ of eq. (11) has a variety of properties. The proofs of all results presented in this section can be found in Section B of the Supplementary Materials.

#### Condition 1.

*The functions u, v and w*_*j*_ *of* *eq*. (1) *are differentiable everywhere for all j* ∈ {1, *…, m*}.

#### Proposition 1.

*Under Condition 1, the surrogate* Φ^*μ*^ *is differentiable everywhere with explicit gradient*

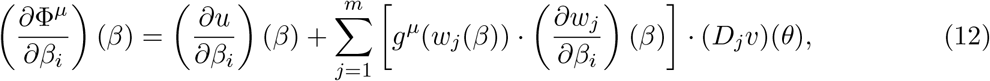

*where g*^*μ*^ *is the derivative of f*^*μ*^ *as given in* *eq*. (7) *and* *eq*. (9), *D*_*j*_*v is the derivative of v with respect to its jth argument, and θ is given by the vector θ* = (*f*^*μ*^(*w*_1_(*β*)), *…, f*^*μ*^(*w*_*m*_(*β*))) ∈ ℝ^*m*^.

The proof of Proposition 1 follows from a direct calculation. Subject to the following commonly satisfied condition, the surrogate Φ^*μ*^ is strictly convex.

#### Condition 2.

*In* *eq*. (1), *the function u is convex and v preserves strict convexity of its input arguments*.

#### Proposition 2.

*Under Condition 2*, Φ^*μ*^ *is strictly convex*.

The proof of Proposition 2 follows from the fact that according to Proposition 6 in Section B, 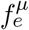 and 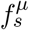 are strictly convex. Thus if *u* is convex and *v* preserves strict convexity, Φ^*μ*^ is itself strictly convex as the sum of a convex function and a strictly convex function. Alternatively, if *v* only preserves convexity, Φ^*μ*^ will still be convex (given *u* is convex). However, the results of Propositions 4 and 5 below do not hold true any more as they require strict convexity.

Next, we consider an approximation bound of Φ^*μ*^ to Φ. Under the following condition, the bounds on 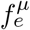 in eq. (8) and 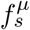 in eq. (10) translate to a guaranteed error bound on the surrogate Φ^*μ*^.

#### Condition 3.

*The function v of* *eq*. (1) *is Lipschitz continuous with Lipschitz constant L*_*v*_.

Condition 3 can easily be verified. Given Condition 1 is already satisfied, and *v* is thus differentiable, then a bounded derivative (in *L*_1_ norm) of *v* will make *v* Lipschitz continuous with 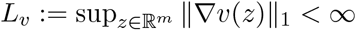.

Condition 3 is used in the following proposition to establish error bounds for Φ^*μ*^ with respect to Φ.

#### Proposition 3.

*Let C*^*μ*^ *denote either* 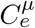 *of* *eq*. (8) *or* 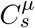 *of* *eq*. (10) *depending on whether* 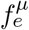 *or* 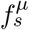 *are employed in* Φ^*μ*^. *Under Condition 3*,

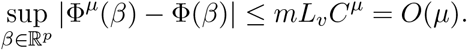

In Proposition 3, the dimension *m* (which oftentimes is taken to be *p*, see Section 2.4) and the Lipschitz constant *L*_*v*_ are fixed for a particular estimation problem, thus allowing us to make the approximation error arbitrarily small as the smoothing parameter *μ* → 0.

Importantly, we are not only interested in the error bound between the objective function Φ and its surrogate Φ^*μ*^, but in how the minimizer 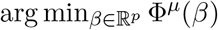 compares to the one obtained had we computed 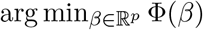.

Following Lemma 2.9 (Seijo & Sen, 2011), the following proposition shows that continuity, strict convexity and a vanishing approximation error of the surrogate implies that the global minimizers of Φ and Φ^*μ*^ converge to each other in the supremum norm metric (again assuming *m* is kept fixed).

#### Proposition 4.

*Let F*_1_ : ℝ^*s*^ → ℝ *be continuous and strictly convex for s* ∈ ℕ. *Then* 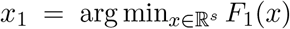 *is continuous at F*_1_ *with respect to the supremum norm*.

Under Conditions 1–3, Φ^*μ*^ satisfies the requirements of Proposition 4. Using the result of Proposition 3, 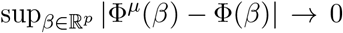 for *μ* → 0 implies that the minimizers of Φ^*μ*^ and Φ converge to each other in the supremum norm. This result is stronger than the one of Theorem 4.4 in the original publication of the FISTA algorithm(Beck & Teboulle, 2009), where it is proven that the FISTA method finds a minimizer which is of similar quality than the true minimizer.

Although Proposition 4 shows convergence, it does not give an explicit error bound on the distance between the two minimizers. This is done in the next result.

#### Proposition 5.

*Let s* ∈ ℕ *and ϵ >* 0. *Let F*_1_ : ℝ^*s*^ → ℝ *be differentiable and strictly convex. Let F*_2_ : ℝ^*s*^ → ℝ *be such that* 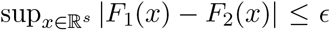. *Let* 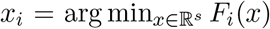 *be the two minimizers for i* ∈ {1, 2}. *Then for any δ >* 0, *there exist two constants C*_*δ*_ *>* 0 *and L*_*δ*_ *>* 0 *independent of x*_2_ *such that*

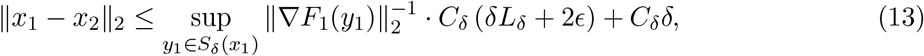

*where S*_*δ*_(*x*) = {*y* : ∥*x* − *y* ∥ _2_ = *δ*} ⊂ ℝ^*s*^ *denotes the sphere of radius δ around x* ∈ ℝ^*s*^ *with respect to the Euclidean norm*.

Note that Proposition 5 does not generalize to non strictly convex functions. Importantly, 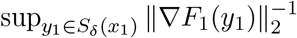 in eq. (13) is well-defined since it suffices to take the supremum over the sphere of radius *δ* around *x*_1_, not the ball of radius *δ* around *x*_1_, thus avoiding the case that ∥∇*F*_1_(*y*_1_) ∥_2_ = 0 at the minimizer *y*_1_ = *x*_1_.

Under Conditions 1–3, applying Proposition 5 with *F*_1_ taken to be the differentiable and strictly convex Φ^*μ*^ and *F*_2_ taken to be Φ immediately gives an explicit bound on the distance between the two minimizers 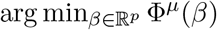 and 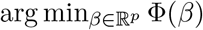. This bound can be computed using the minimizer of Φ^*μ*^ only. We demonstrate the quality of the bound in the simulations of Section 3.3.

### 2.4 Applications to popular L1 penalized regression operators

Let *X* ∈ ℝ^*n*×*p*^, *y* ∈ ℝ^*n*^, and *β* ∈ ℝ^*p*^ as introduced in Section 1. Additionally, let *λ, α, γ* ≥ 0. A variety of popular regression operators can be written in the form of eq. (1) and satisfy the conditions of Section 2.3.

1. The least absolute shrinkage and selection operator (Lasso) (Tibshirani, 1996) is defined as

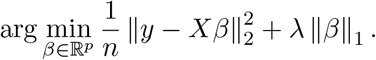 It can be expressed in the form of eq. (1) with 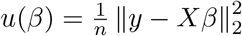, as well as 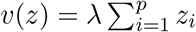 for *z* ∈ ℝ^*p*^, and *w*_*j*_(*β*) = *β*_*j*_ for *j* ∈ {1, *…, p*}. Here, 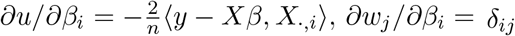, and *D*_*j*_*v* = *λ* for *i, j* ∈ {1, *…, p*}.
2. The elastic net (Zou & Hastie, 2005) augments the Lasso with an *L*_2_ penalty on the parameters *β*, resulting in

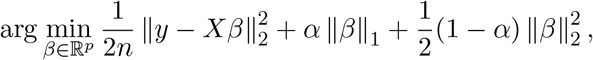

using a published parameterization (Friedman et al., 2010b) which is also used in the specification of the R-package *glmnet* (Friedman et al., 2020). It can be expressed in the form of eq. (1) with 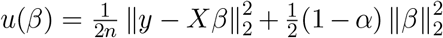, as well as 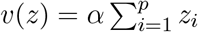 for *z* ∈ ℝ^*p*^ and *w*_*j*_ as in the previous case of the standard Lasso, where *j* ∈ {1, *…, p*}. The derivative of *u* is 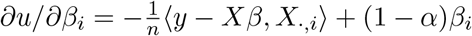
3. The fused Lasso (Tibshirani et al., 2005) is defined as

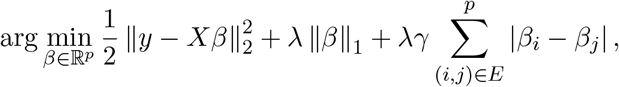

where *E* is the edge set of some underlying graph (Arnold & Tibshirani, 2020). Let *E* = {*e*_1_, *…, e* |_*E*_ |) in some arbitrary ordering, and 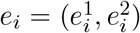 for all *i* ∈ {1, *…*, |*E*|}. The fused Lasso can be casted in the form of eq. (1) with *u*(*β*) defined similarly to the standard Lasso, as well as 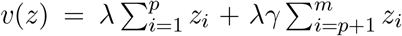 for *m* = *p* + |*E*| and *z* ∈ ℝ^*m*^. Accordingly, *w*_*j*_(*β*) = *β*_*j*_ for *j* ∈ {1, *…, p*} and 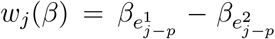 for *j* ∈ {*p* + 1, *…, m*}. Here, *D*_*j*_*v* = *λ* for *j* ∈ {1, *…, p*} and *D*_*j*_*v* = *λγ* for *j* ∈ {*p* + 1, *…, m*}. Moreover, *∂w*_*j*_/*∂β*_*i*_ = *δ*_*ij*_ for *j* ∈ {1, *…, p*} and 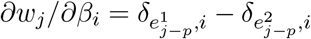 for *j* ∈ {*p* + 1, *…, m*}.
4. The graphical Lasso (Friedman et al., 2008, 2019) considers observations *X*_1_, *…, X*_*n*_ ∼*N* (0, Σ) from a multivariate Gaussian distribution and estimates the precision matrix Θ = Σ^−1^ ∈ ℝ^*p*×*p*^ as

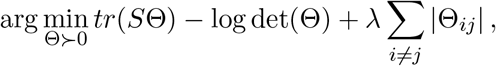

where *S* is the sample covariance matrix, *tr* denotes the trace of a matrix, and the arg min is taken over all positive definite matrices Θ. Define 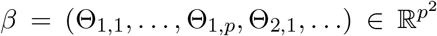 as the vector containing all parameters of Θ stacked together by row. As before, we define the differentiable part of eq. (1) to include the trace and log terms, thus *u*(Θ) = *tr*(*S*Θ) − log det(Θ) with derivative ∇*u* = *S*^T^ − Θ^−1^, and *v* : ℝ^*m*^ → ℝ and 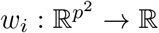 with *m* = *p*^2^ as in the case of the standard Lasso.

#### Algorithm 1: Progressive smoothing

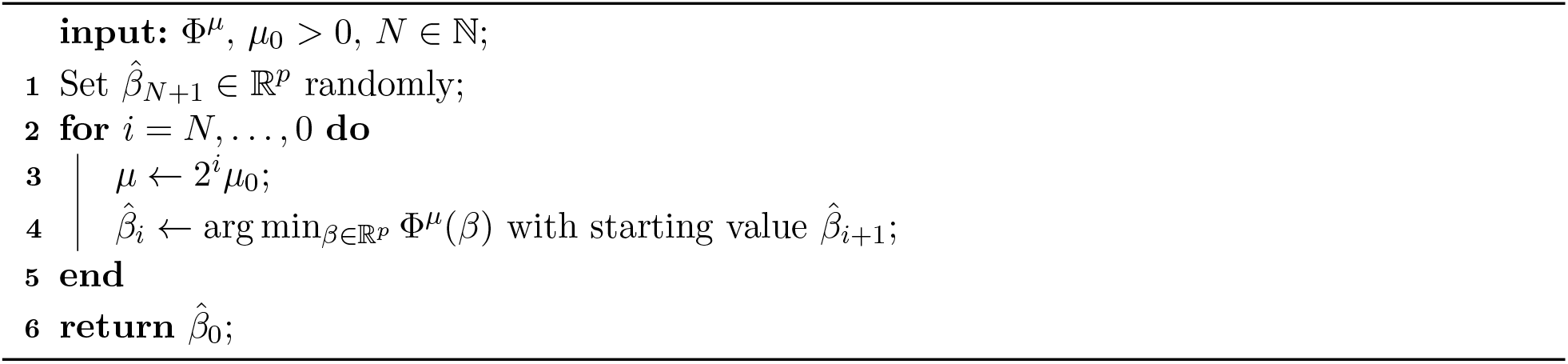

In all above cases, the functions *u, v*, and *w*_*i*_ are differentiable for *i* ∈ {1, *…, m*}, thus satisfying Condition 1. Their derivatives are given explicitly for convenience.

Moreover, all functions *u* above, including the one of the graphical Lasso, are convex as they can be written as the linear combinations of (Euclidean) norms. Importantly, the functions *v* are always linear combinations of their inputs with positive coefficients and thus preserve strict convexity. Therefore, Condition 2 is satisfied.

Finally, the functions *v* are all differentiable and have a bounded gradient (in *L*_1_ norm), thus making them Lipschitz continuous as required in Condition 3. For instance, 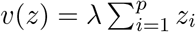 for *z* ∈ ℝ^*p*^ in case of the standard Lasso, thus we obtain ∇*v* = *λ*[1, *…*, 1] and 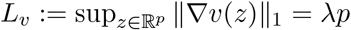 using the fact that *λ* is non-negative.

### 2.5 Progressive smoothing algorithm

This section proposes an adaptive smoothing technique for the surrogate Φ^*μ*^. Simulations in Section 3 show that the progressive smoothing algorithm yields stable estimators for linear regression.

Instead of solving the smoothed problem 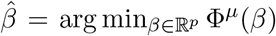 directly for some *μ >* 0, we employ a progressive smoothing procedure along the following rationale: We start with a large value of the smoothing parameter *μ* to facilitate the minimization. After computing 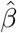, we decrease the smoothing parameter and repeat the minimization using the previously found minimizer as the new starting value. This approach is based on the heuristic idea that as *μ* decreases and the smoothed surrogate Φ^*μ*^ approaches Φ, the found minimizers in each iteration remain close to each other and converge to the minimizer of Φ.

Algorithm 1 formalizes our approach. The input of the algorithm is the function Φ^*μ*^, a target smoothing parameter *μ*_0_ *>* 0, and a number of smoothing steps *N* ∈ ℕ. After initializing a random starting value 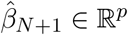 for the first minimization, we gradually decrease the degree of smoothness according to *μ* = 2^*i*^*μ*_0_ from *i* = *N* to the target level *μ*_0_ at *i* = 0. In each iteration *i*, we compute a new estimate 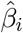using the current smoothing level *μ* and the previous estimate 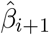 as the starting value. The output of the algorithm is 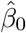, the parameter estimate corresponding to the target smoothing degree *μ*_0_.

Importantly, the advantage of Algorithm 1 consists in the fact that the precise specification of the smoothing parameter does not play a major role. It suffices to start with any sufficiently large value (that is, 2^*N*^ *μ*_0_ ≫ 1) and to end the iteration with any sufficiently small value *μ*_0_, for instance of the order of the machine precision or of the square root of the machine precision. This effectively makes Algorithm 1 free of tuning parameters.

## 3 Experimental results

In this section, we evaluate the performance of the regression estimates computed with our proposed smooth surrogate Φ^*μ*^ of eq. (11). We benchmark against the four approaches considered in Section 2.4. We start with results for the standard Lasso on simulated data (Section 3.1), followed by the elastic net (Section 3.2). Further experiments on simulated data with the fused Lasso and the graphical Lasso are reported in Section C of the Supporting Information. The quality of the theoretical bound stated in Proposition 5 is assessed in Section 3.3.

We implement the three methodologies we developed in this paper:

1. We carry out the minimization of the unsmoothed Φ of eq. (1) using R’s function *optim*. Within the *optim* function, we select the quasi-Newton method *BFGS* for which we supply its explicit (though non-smooth) gradient

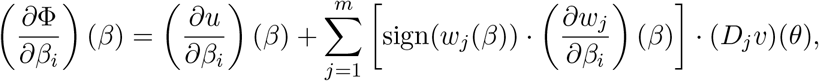

where the sign function is defined as sign(*z*) = 1 for *z >* 0, sign(*z*) = −1 for *z <* 1, and zero otherwise. This approach will be referred to as *unsmoothed Phi*.
2. We minimize the smooth surrogate Φ^*μ*^ of eq. (11) using its explicit gradient given in eq. (12) based on the entropy prox function of Section 2.2. The smoothing parameter of the smooth surrogate Φ^*μ*^ was fixed at *μ* = 0.1. This approach will be referred to as *smoothed Phi*.
3. We employ the progressive smoothing approach of Section 2.5. As suggested at the end of Section 2.5, we set the target smoothing parameter to *μ*_0_ = 2^−6^ (roughly the square root of the machine precision) and employ *N* = 9 smoothing steps (thus implying an initial value of the smoothing parameter of *μ* = 2^−6^ · 2^9^ = 2^3^).

We benchmark our methods against the gold standards available in the literature for the four problems we consider:

1. As a general method, we use the Fista algorithm to minimize Φ as implemented in the *fasta* R-package (Chi et al., 2018). The main function of the *fasta* package which implements the Fista algorithm, also called *fasta*, requires the separate specification of the smooth and non-smooth parts of the objective function including their explicit gradients. We follow Example 2 in the *vignette* of the *fasta* R-package (Chi et al., 2018) and supply both as specified in eq. (1). Moreover, in contrast to our approach, a variety of tuning parameters need to be selected by the user, e.g. an initial starting value, an initial stepsize, parameters determining the lookback window for non-monotone line search and a shrinkage parameter for the stepsize. The initial stepsize for Fista is set to *τ* = 10 as in Example 2 of the package vignette (Chi et al., 2018). The lookback window for non-monotone line search and the shrinkage parameter for the stepsize are left at their default values. We employ a uniformly random starting value for *β* ∈ ℝ^*p*^ (each vector entry is drawn independently from *U* [0, 1]) as done for our own approaches.
2. For the elastic net comparison, we benchmark against the *glmnet* algorithm (Friedman et al., 2010a), available in the R-package *glmnet* (Friedman et al., 2020) on CRAN. The glmnet algorithm is a variant of Fista which performs a cyclic update of all coordinates, whereas Fista updates all coordinates per iteration. To solve an elastic net problem with glmnet, we set its parameter *alpha* to 0.5 as specified on page 24 of the package vignette (Friedman et al., 2020).
3. For the fused Lasso, we employ the R-package *genlasso* (Arnold & Tibshirani, 2020) on CRAN with default parameters. The fused Lasso is implemented in the function *fusedlasso*.
4. For the graphical Lasso, we employ the R-package *glasso* (Friedman et al., 2019) on CRAN with default parameters. The function to run the graphical Lasso is likewise called *glasso*.

Two algorithms are unconsidered in this article for the following reasons. Since the R-package *SIS* (Fan & Li, 2001) does itself rely on glmnet for computing regression estimates, we omit it in this section. The LARS algorithm (Efron et al., 2004) is implemented in the R-package *lars* (Hastie & Efron, 2013) on CRAN. As remarked in the literature (Friedman et al., 2010a), LARS is slower than glmnet or Fista. Additionally, since the LARS implementation (Hastie & Efron, 2013) always computes a full LASSO path, it is considerably slower than the aforementioned methods.

All results are averages over 100 repetitions. The regularization parameters of Section 2.4 were fixed at *λ* = 1, *α* = 0.5, and *γ* = 0.5. We evaluate the accuracy of the aforementioned algorithms using two metrics. First, we report the *L*_2_ norm of the true parameters *β* that we generate to the estimated ones returned by each method. Second, we report the standard vector correlation between true and estimated parameters. To assess computational efficiency and runtime scaling, we report log-log plots of the empirical runtimes we measure.

### 3.1 Standard Lasso on simulated data

We start by applying our smoothing framework to the standard Lasso (Tibshirani, 1996). To this end, we employ the choices of *u, v*, and *w*_*j*_, *j* ∈ {1, *…, p*}, for the standard Lasso given in Section 2.4.

We simulate a regression model of the type *y* = *Xβ* by drawing the rows of the design matrix *X* ∈ ℝ^*n*×*p*^ from a multidimensional normal distribution with the following mean vector and covariance matrix. The entries of the *p* dimensional mean vector of the multidimensional normal distribution are drawn independently from a uniform distribution in [0, 1]. To ensure positive definiteness, the *p* × *p* dimensional covariance matrix of the multidimensional normal distribution is drawn from a Wishart distribution with sample size 1, *p* degrees of freedom, and scale matrix set to the *p* × *p* dimensional identity matrix.

After generating *X*, we draw the true parameters *β* independently from a standard normal distribution. To ensure sparseness, we set all but *nz* ∈ {0, *…, p*} out of the *p* entries to zero. We fix *nz* = 0.2*p* in the entire simulation section. Finally, we calculate *y* ∈ ℝ^*n*^ as *y* = *Xβ* + *ϵ* for some noise vector *ϵ* ∈ ℝ^*n*^. The entries of *ϵ* are generated independently from a Normal distribution with mean zero and some variance *σ*^2^. The smaller we choose *σ*^2^, the easier the recovery of *β* will be. We fix *σ*^2^ = 0.1.

We benchmark our three approaches (unsmoothed Φ, smoothed surrogate Φ^*μ*^, and progressive smoothing) against the Fista algorithm. We are interested in accuracy and runtime of all algorithms as a function of *n* (while keeping *p* = 100 fixed) or *p* (while keeping *n* = 100 fixed).

Figure 1 shows results for the scaling in *n* while keeping *p* = 100 fixed. We observe that, as expected, estimation becomes easier as *n* becomes larger. Fista yields the best estimates when measuring the error in the *L*_2_ norm, followed by the unsmoothed Lasso objective (unsmoothed Phi) and the progressive smoothing algorithm. Minimizing the smoothed Lasso objective (smoothed Phi) does not seem to have an advantage in this scenario. The correlation of the truth to the estimate seems equally high for almost all approaches. Moreover, we observe that all approaches have roughly the same runtime scaling in *n*. Since the progressive smoothing approach works by repeatedly minimizing the smoothed surrogate, its runtime is roughly a constant factor higher than the one for minimizing the smoothed surrogate Φ^*μ*^.

**Figure 1:**
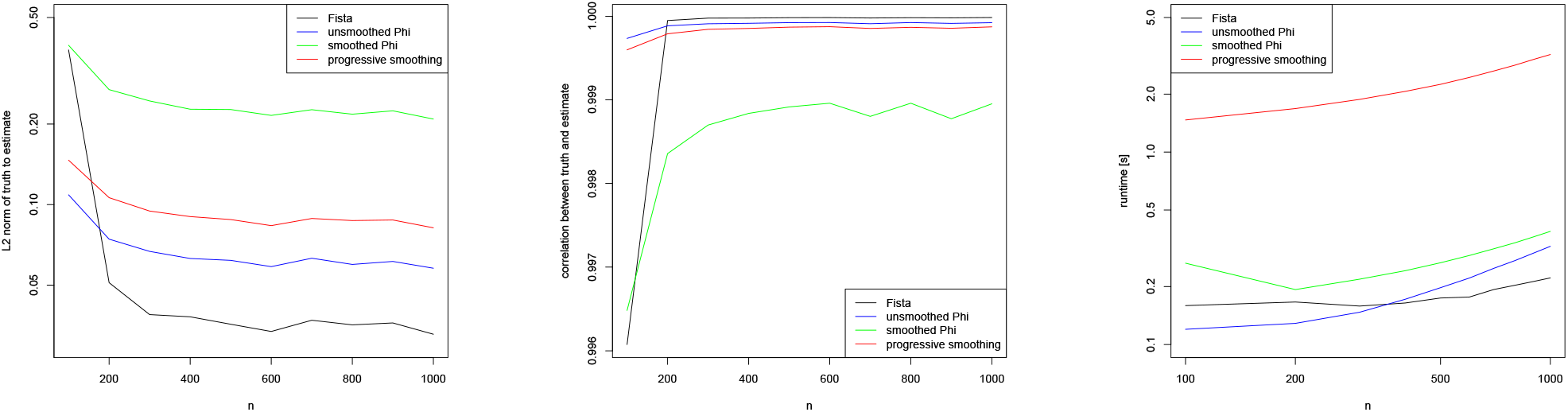
Standard Lasso: *L*_2_ norm (left) and correlation (middle) between simulated truth and parameter estimates as a function of *n* while *p* = 100. Runtime in seconds (right) has a log scale on both axes. A color version of this figure can be found in the electronic version of the article.

The more important case is the scenario in which *p* ≫ *n*. This scenario, depicted in Figure 2, shows a different picture. Fista does not seem to approximate the truth well when measuring the error with the *L*_2_ norm, whereas minimizing the smoothed surrogate (smoothed Phi) and progressive smoothing yield much more accurate results. The picture is confirmed when measuring accuracy with the correlation between truth and estimate. Here, using Fista and minimizing the unsmoothed objective function results in estimates with a considerably lower correlation to the true parameter vector than using both our smoothing approaches (smoothed Phi or progressive smoothing). As can be seen from the left and middle panels in Figure 2, minimizing the unsmoothed Lasso objective Φ is more prone to instabilities due to its non-differentiability. This is to be expected and motivates our smoothing approach. Not surprisingly, while keeping the data size *n* fixed, the estimation becomes more challenging as *p* increases. The runtime scaling of our three methods is again similar, though Fista seems to have a slightly more favorable runtime scaling in *p*.

**Figure 2:**
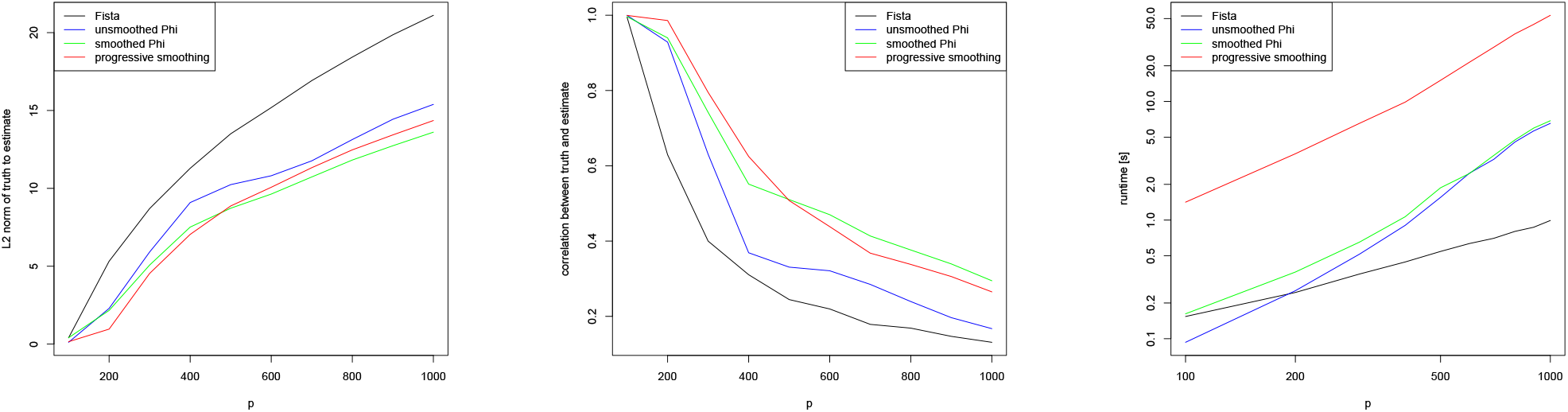
Standard Lasso: *L*_2_ norm (left) and correlation (middle) between simulated truth and parameter estimates as a function of *p* while *n* = 100. Runtime in seconds (right) has a log scale on both axes. A color version of this figure can be found in the electronic version of the article.

### 3.2 Elastic net on simulated data

We repeat the experiments of Section 3.1 for the elastic net (Zou & Hastie, 2005) using the R-package *glmnet* (Friedman et al., 2010a). For this, we employ the choices of *u, v*, and *w*_*j*_, *j* ∈ {1, *…, p*}, for the elastic net given in Section 2.4. We generate models *y* = *Xβ* as described in Section 3.1, and evaluate Fista, glmnet, unsmoothed Φ, smoothed surrogate Φ^*μ*^, and progressive smoothing with respect to their accuracy and scaling behavior in *n* and *p*.

Figure 3 shows scaling results for *n* while keeping *p* = 100 fixed. We observe that glmnet and the simple smoothing approach (smoothed Phi) do not seem to perform well in this experiment, while Fista as well as the minimization of the unsmoothed objective Φ and progressive smoothing perform notably better. When looking at the correlation between truth and estimate, all methods with the exception of glmnet and the smoothed surrogate Φ^*μ*^ (smoothed Phi) again perform similarly. The runtime of all methods does not seem very dependent on *n*.

**Figure 3:**
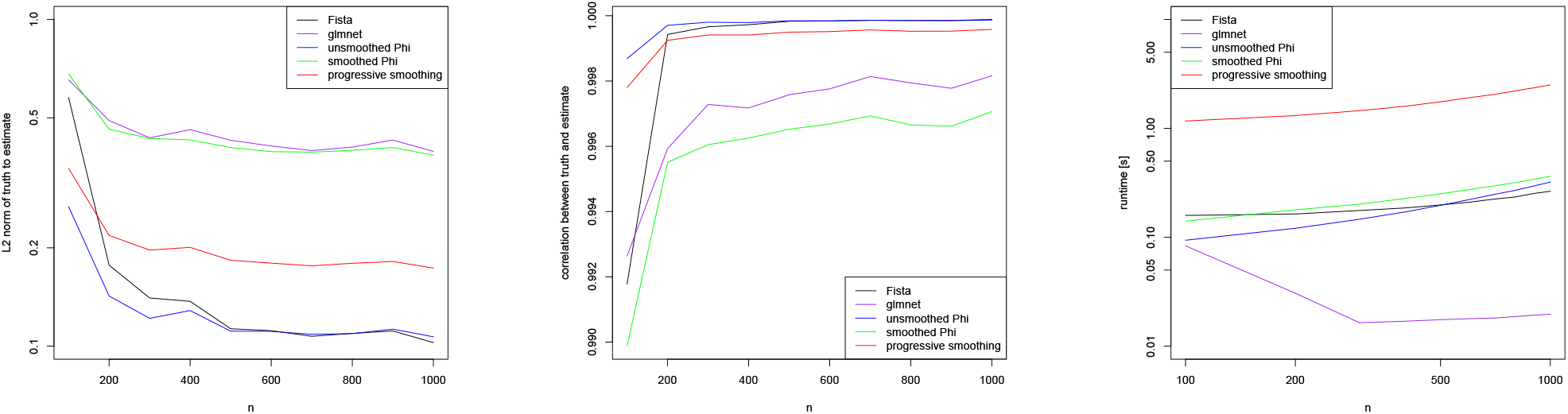
Elastic net: *L*_2_ norm (left) and correlation (middle) between simulated truth and parameter estimates as a function of *n* while *p* = 100. Runtime in seconds (right) has a log scale on both axes. The comparison includes Fista, glmnet, unsmoothed Φ, smoothed surrogate Φ^*μ*^, and progressive smoothing.

Analogously, Figure 4 presents scaling results for the more interesting case that *p* ≫ *n* while keeping *n* = 100 fixed. These results confirm the observations made for the case of the standard Lasso in Figure 2. As expected, the estimation of the parameters becomes more challenging for all methods as *p* increases while *n* stays fixed.

**Figure 4:**
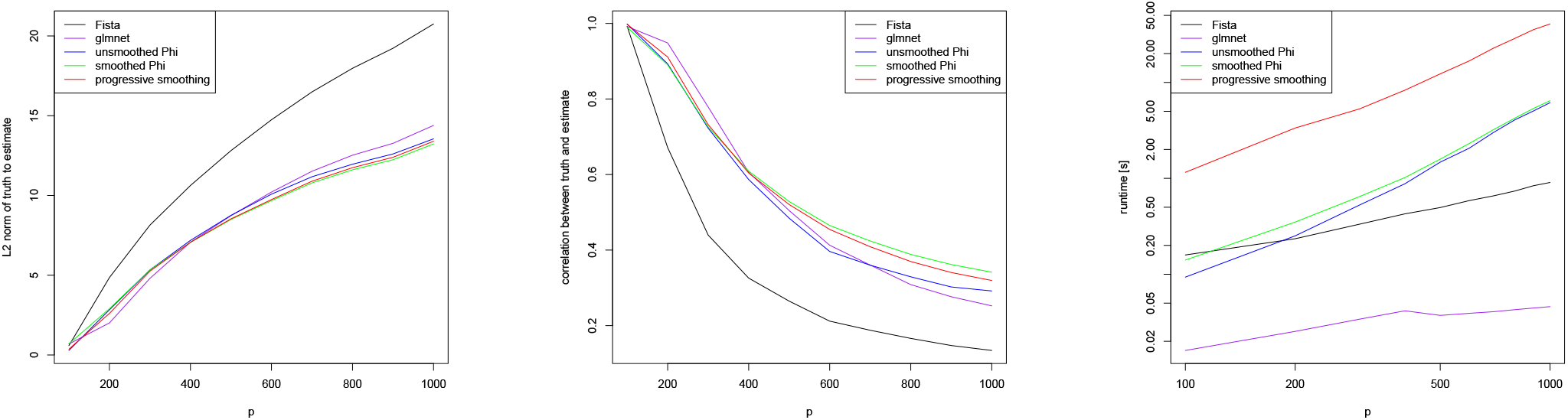
Elastic net: *L*_2_ norm (left) and correlation (middle) between simulated truth and parameter estimates as a function of *p* while *n* = 100. Runtime in seconds (right) has a log scale on both axes. The comparison includes Fista, glmnet, unsmoothed Φ, smoothed surrogate Φ^*μ*^, and progressive smoothing.

Fista seems to yield estimates of poorest quality when assessing accuracy with the *L*_2_ distance between truth and estimate, while the other approaches perform more favorably. Regarding the correlation between truth and estimate, Fista again performs suboptimally, while glmnet and the minimization of the unsmoothed Φ (unsmoothed Phi) perform better. Using our smoothed surrogate (smoothed Phi) and the progressive smoothing algorithm perform best in that they retain the largest correlation between their estimates to the true parameters. As expected, the runtime scaling in *p* of Fista and glmnet is roughly similar, and the asymptotic runtimes of both seem to be lower than for our methods. Minimizing the unsmoothed objective function and the smooth surrogate seems to have an almost identical asymptotic runtime, while progressive smoothing is a constant factor slower as expected.

### 3.3 Verification of theoretical bounds

Finally, we verify the theoretical bounds derived in Proposition 5 for the case of the standard Lasso using simulated data. In this experiment, we therefore again employ the choices of *u, v*, and *w*_*j*_, *j* ∈ {1, *…, p*}, for the standard Lasso given in Section 2.4.

First, we compute regression estimates with the unsmoothed and smoothed Lasso objective functions as done in Section 3.1. We employ a regularization parameter of *λ* = 1, and a smoothing parameter of *μ* = 1. The generation of the model *y* = *Xβ* was done as described in Section 3.1.

After performing the minimization we record ∥ *x*_1_ − *x*_2_ ∥_2_, the left hand side of eq. (13). This quantity is assumed to be unknown as we aim to avoid the computation of the minimizer *x*_2_ of the unsmoothed objective and solely focus on the minimizer *x*_1_ of the smoothed surrogate instead.

Moreover, we compute the right hand side of eq. (13), given by the expression 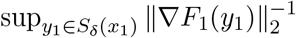. *C*_*δ*_ (*δL*_*δ*_ + 2*ϵ*) + *C*_*δ*_*δ* In the case of the standard Lasso considered here, *F*_1_ is the differentiable and strictly convex function Φ^*μ*^, that is the smooth surrogate of the standard Lasso (see the discussion at the end of Section 2.3). The parameter *δ* is chosen by the user: for more stable results, we compute the bound of eq. (13) on the grid *δ* ∈ {0.1, 0.2, *…*, 1} and take the minimal value. The value of is given by eq. (8). Taking the supremum over the parameter *y*_1_ on the sphere *S*_*δ*_(*x*_1_) is performed in a Monte Carlo fashion by computing the bound of eq. (13) for random points generated uniformly on *S*_*δ*_(*x*_1_) and taking the maximum. Last, *C*_*δ*_ can be computed explicitly using a scalar product between two known quantities as shown in the proof of Proposition 5, and *L*_*δ*_ is taken to be the local Lipschitz constant of Φ^*μ*^ at its minimizer *x*_1_. This makes the right hand side of eq. (13) explicitly computable without knowledge of the minimizer *x*_2_ of the unsmoothed (original) Lasso objective.

Results for the left and right hand sides of eq. (13) are shown in Figure 5, once for the scaling in *n* while *p* = 100 is kept fixed, and once for the scaling in *p* while *n* = 100 is kept fixed. As expected, the estimation becomes easier as *n* increases for a fixed *p*, and more challenging as *p* increases for a fixed *n*.

**Figure 5:**
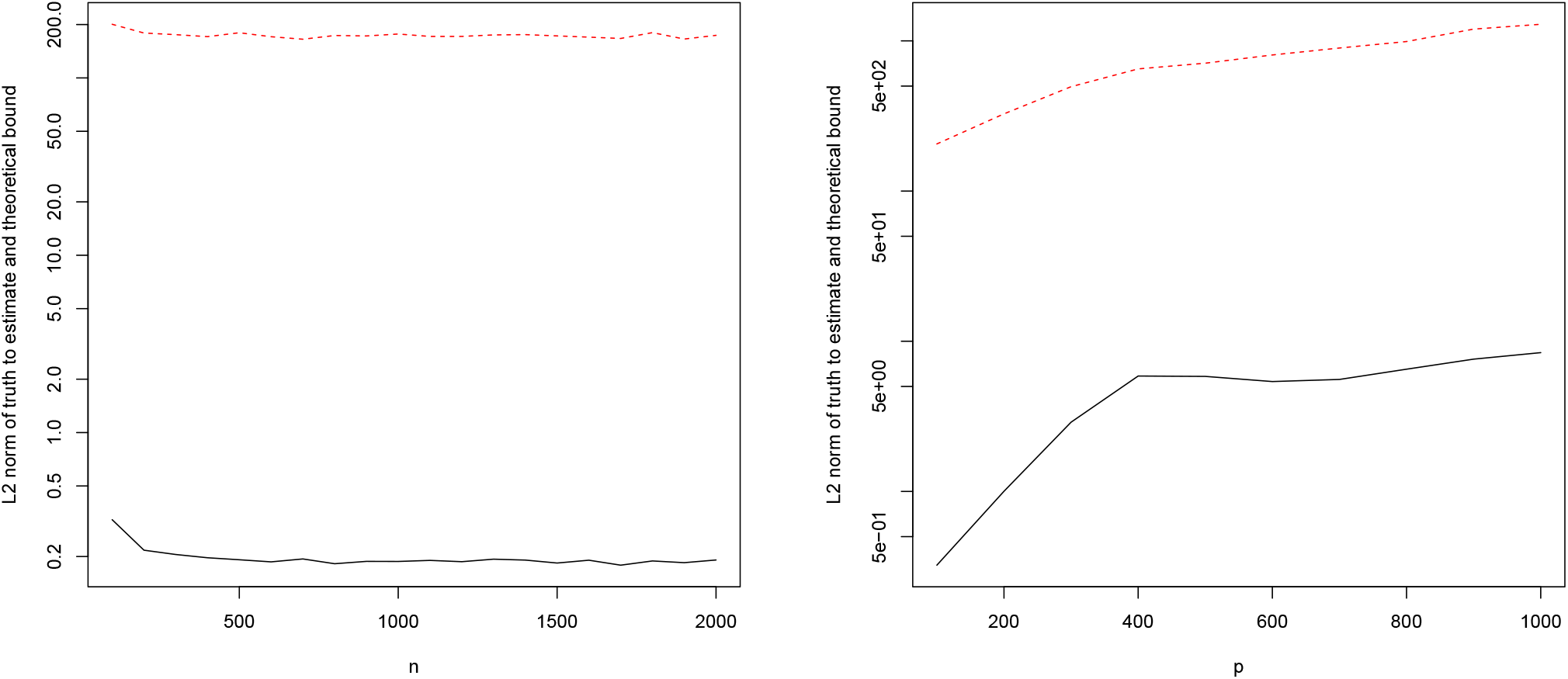
True *L*_2_ norm between minimizers of unsmoothed and smoothed Lasso (left hand side of eq. (13)) in black, and theoretical upper bound (right hand side of eq. (13)) in red. Scaling in *n* while *p* = 100 is fixed (left) and scaling in *p* while *n* = 100 is fixed (right). Log scale on the y-axis. A color version of this figure can be found in the electronic version of the article.

Overall, we observe that eq. (13) indeed provides a valid, explicit upper bound on the error between the unsmoothed and smoothed minimizers for each fixed choice of *μ*. For the scaling in *n*, the bound is very conservative, but in the more challenging scenario of the scaling in *p*, the bound seems to accurately follow the error we make by using the smoothed surrogate. Note that by Propositions 3 and 5, this error will eventually go to zero as the smoothing parameter *μ* → 0.

## 4 Discussion

In this paper, we propose a smoothing framework for a general class of *L*_1_ penalized regression operators that includes most common approaches, e.g. the Lasso, elastic net, fused and graphical Lasso, etc. Throughout the article, we consider a fixed parameter setting (parameters *n, m, p* are fixed, see Section 1). The framework is based on a closed-form smooth surrogate and its closed-form gradient. Since the aforementioned regression operators are convex but non-smooth, due to the non-differentiability of their *L*_1_ norm, minimizing the smooth surrogates instead allows for a fast and efficient computation of regression estimates.

Most importantly, we prove a series of theoretical results which apply to the entire class of regression operators belonging to our framework. Under regularity conditions, we show that the smooth surrogate is both strictly convex and uniformly close (in supremum norm) to the original (unsmoothed) objective function. This has the important implication that the regression estimates obtained by minimizing the smooth surrogate can be made arbitrarily close to the ones of the original (unsmoothed) objective function. Additionally, we provide explicitly computable error bounds on the accuracy of the estimates obtained with the smooth surrogate. Since we can efficiently carry out the minimization of the smooth surrogate, our approach yields easily computable regression estimates that are both guaranteed to be close to the estimates of the original objective function, and allow for a priori error estimation. Furthermore, we develop a progressive smoothing algorithm for iterative smoothing. This approach is virtually free of tuning parameters. Its runtime is only a constant factor higher than that of the other algorithms (due to repeated optimization calls).

Smoothing the *L*_1_ penalty of the aforementioned regression operators typically causes the resulting regression estimates to be dense. Although the regression estimates for those variables which are shrunk to zero by the original regression operators (with *L*_1_ penalty) will typically be very small, smoothing the *L*_1_ penalty in our framework thus somewhat leads to a loss of the variable selection property. However, an easy fix to re-establish sparse estimates and enable variable selection is to apply some form of thresholding.

Our experimental results support our theoretical conclusions that minimizing the proposed smooth surrogate yields estimates of equal or better quality than many of the gold standard approaches available in the literature (the Fista, glmnet, gLasso algorithms, etc.). At the same time, our approach has roughly the same runtime scaling as the established approaches we include in our comparisons while keeping all aforementioned theoretical guarantees.

## Funding

The initial methodology work for this paper was funded by Cure Alzheimer’s Fund; Funding for this research was provided through the National Heart, Lung, and Blood Institute [U01HL089856, U01HL089897, P01HL120839, P01HL132825, 2U01HG008685] and National Human Genome Research Institute [R01HG008976].

## Supporting Information

The Supporting Information contains a detailed overview of Nesterov smoothing (Section A), all proofs (Section B), as well as additional simulations (Section C).

## Supporting Information

### A Nesterov smoothing

This section follows Sections 2 and 4 of Nesterov’s published work (Nesterov, 2005). It introduces the basic formalism of Nesterov smoothing in Section A.1 and concretizes the approach in Section A.2.

#### A.1 Description of Nesterov smoothing

We are given a piecewise affine and convex function *f* : ℝ^*q*^ → ℝ which we aim to smooth, where *q* ∈ ℕ. We assume that *f* is composed of *k* ∈ ℕ linear pieces (components). The function *f* can be expressed as

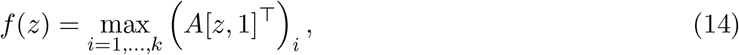

where *A* ∈ ℝ^*k*×(*q*+1)^ is a matrix whose rows contain the linear coefficients for each of the *k* pieces (with the constant coefficients being in column *q* + 1), *z* ∈ ℝ^*q*^, and [*z*, 1] ∈ ℝ^*q*+1^ denotes the vector obtained by concatenating *z* and the scalar 1 (see Section 2).

Let ∥ · ∥^*k*^ be a norm on ℝ^*k*^ and ⟨ ·, · ⟩ be the Euclidean inner product. Define the unit simplex *Q*_*k*_ ⊆ ℝ^*k*^

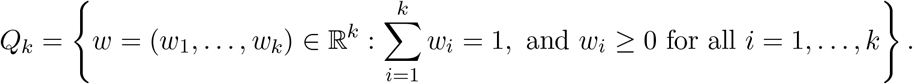

To introduce the smoothing procedure, a proximity function (or prox function) is defined first on *Q*_*k*_ (Nesterov, 2005). A prox function *ρ* is any nonnegative, continuously differentiable, and strongly convex function (with parameter *σ >* 0 and with respect to the norm ∥ · ∥ _*k*_). The latter means that *ρ* satisfies

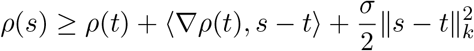

for all *s, t* ∈ *Q*_*k*_.

For any *μ >* 0, consider the function

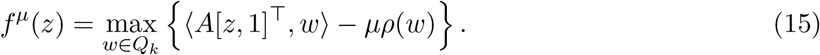

According to Theorem 1 in Nesterov’s original publication (Nesterov, 2005), the function *f*^*μ*^ defined in eq. (15) is convex and everywhere differentiable in *z* for any *μ >* 0. The function *f*^*μ*^ depends only on the parameter *μ* controlling the degree of smoothness. For *μ* = 0, we recover the original unsmoothed function since *f* (*z*). The gradient 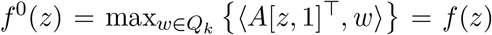. The gradient 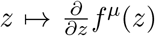 is Lipschitz continuous with a Lipschitz constant that is proportional to *μ*^−1^. A closed form expression of both the gradient and the Lipschitz constant are given in Theorem 1 in Nesterov’s original publication (Nesterov, 2005).

Importantly, the function *f*^*μ*^ is a uniform smooth approximation of *f* = *f* ^0^ since

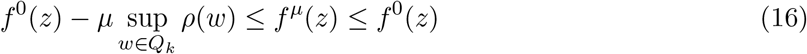

for all *z* ∈ ℝ^*q*^, meaning that the approximation error is uniformly upper bounded by

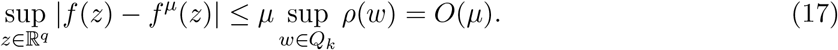

Indeed, eq. (16) holds true since for all *z* ∈ ℝ^*q*^,

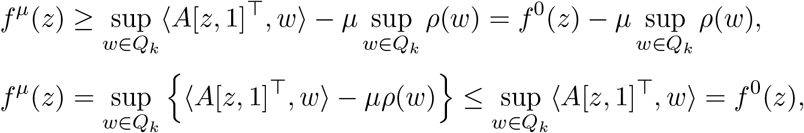

where it was used that both the function *ρ* and the parameter *μ* are nonnegative.

#### A.2 Two choices for the proximity function

We consider two choices of the prox function *ρ*.

##### A2.1. Entropy prox function

The entropy prox function *ρ*_*e*_ : ℝ^*k*^ → ℝ is given by

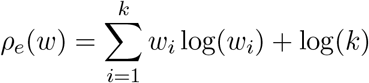

for *w* ∈ ℝ^*k*^.

Setting the norm ∥ · ∥ _*k*_ as the *L*_1_ norm in ℝ^*k*^, Lemma 3 of Nesterov’s original publication (Nesterov, 2005) shows that *ρ*_*e*_ is strongly convex with respect to the *L*_1_ norm and satisfies 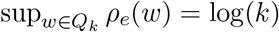 Using eq. (17), we obtain the uniform bound

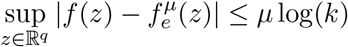

for the entropy smoothed function 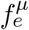 obtained by using *ρ*_*e*_ in eq. (15). Interestingly, smoothing with the entropy prox function admits a closed-form expression of 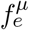 given by

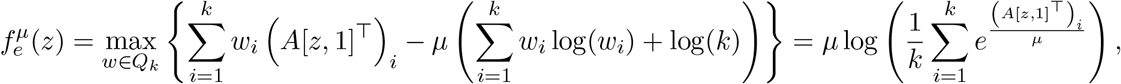

see Lemma 4 of Nesterov’s original publication (Nesterov, 2005).

##### A.2.2 Squared error prox function

The squared error prox function is given by

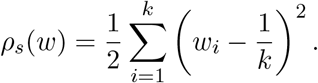

It has been shown (Mazumder et al., 2019) that the optimization in eq. (15) with squared error prox function is equivalent to the convex program

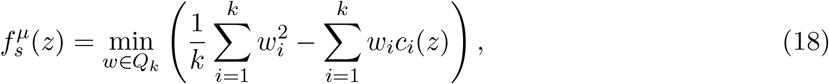

where *c*_*i*_(*z*) = 1*/μ* · (*A*[*z*, 1]^T^)_*i*_ − 1*/k* depends on *A* and *μ* and is defined for any *i* ∈ {1, *…, k*}. The problem in eq. (18) is equivalent to finding the Euclidean projection of the vector *c*(*z*) = (*c*_1_(*z*), *…, c*_*k*_(*z*)) ∈ ℝ^*k*^ onto the *k*-dimensional unit simplex *Q*_*k*_. This projection can be carried out efficiently using the Michelot algorithm (Michelot, 1986), for which a computationally more efficient version was proposed in the literature (Wang & Carreira-Perpiñán, 2013) that we use in our implementations. Denoting the Euclidean projection of the vector *c*(*z*) onto *Q*_*k*_ as vector *ĉ*(*z*), the squared error prox approximation of *f* can be written as

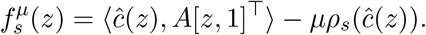

As 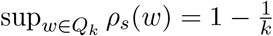 (see Section 4.1 in Nesterov’s original publication Nesterov (2005)), we obtain the uniform bound

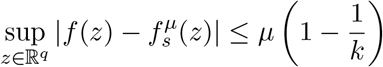

for the squared error smoothing approach.

### B Proofs

#### Proposition 6.

*The entropy prox smoothed function* 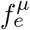 *of* *eq*. (7) *and the squared error prox smoothed* 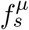 *of* *eq*. (9) *are strictly convex functions*.

#### Proof of Proposition 6.

The second derivative of 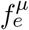 is given by

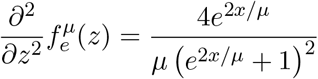

and hence always positive, thus making 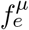 strictly convex. Similar arguments show that 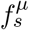 is strictly convex. □

#### Proof of Proposition 3.

Using the fact that according to Condition 3, *v* is Lipschitz continuous with Lipschitz constant *L*_*v*_, we have

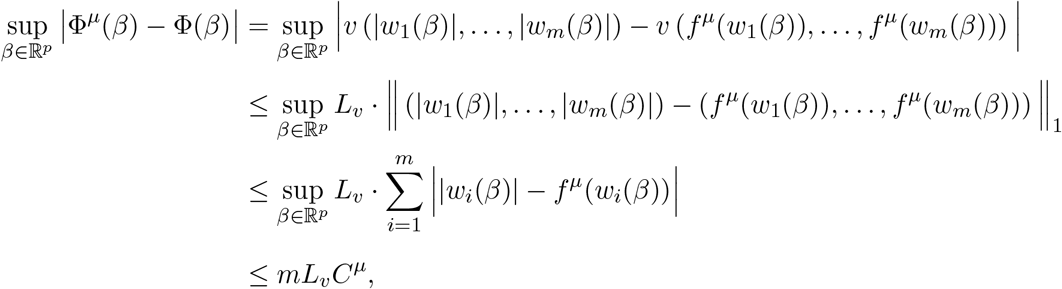

thus establishing the claimed bound. □

#### Proof of Proposition 4.

Since *F*_1_ is continuous and strictly convex, it lays in the Skorohod topology 𝒟_*K*_, see Definition 2.2 (Seijo & Sen, 2011). According to Lemma 2.9 (Seijo & Sen, 2011), the argmax functional is continuous at *F*_1_ with respect to the supremum norm metric. □

#### Proof of Proposition 5.

Let *y*_1_ ∈ *S*_*δ*_(*x*_1_), *y*_1_ ≠ *x*_1_, be such that *x*_1_, *x*_2_ and *y*_1_ are collinear.

Since *F*_1_ is differentiable, we know that ∇*F*_1_ exists. Since *F*_1_ is strictly convex, the minimum *x*_1_ is unique and ∇*F*_1_(*y*_1_) ≠ 0 as *y*_1_ ≠ *x*_1_. Since *F*_1_ is convex, the tangent at every point stays below the function. Thus considering the tangent at *y*_1_, we have for all *z* that

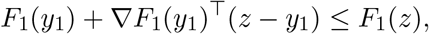

and thus we can bound *F*_1_ − *ϵ* from below as

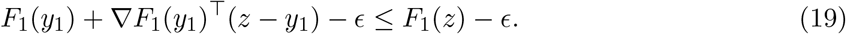

Observe that since 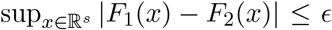, and since *F*_1_ is strictly convex and *F*_2_ is convex, the minimum *x*_2_ of *F*_2_ must satisfy *F*_2_(*x*_2_) ≤ *F*_1_(*x*_1_) + *ϵ*. Moreover, since at *x*_2_ we have *F*_2_(*x*_2_) ∈ [*F*_1_(*x*_2_) − *ϵ, F*_1_(*x*_2_) + *ϵ*], since the tangent of eq. (19) stays below *F*_1_(*x*_2_) − *ϵ*, and since *x*_1_, *x*_2_ and *y*_1_ are collinear, the solution *z*_0_ satisfying

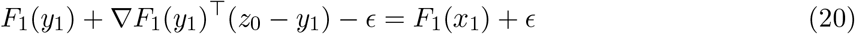

has the property that *x*_2_ cannot be further away from *x*_1_ than *z*_0_, thus ∥*x*_1_ − *x*_2_∥ _2_ ≤ ∥ *x*_1_ − *z*_0_∥ _2_. Rewriting *z*_0_ − *y*_1_ in eq. (20) as *z*_0_ − *x*_1_ + *x*_1_ − *y*_1_ and rearranging terms yields

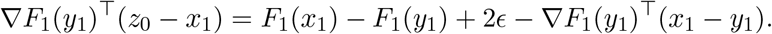

Rewriting ∇*F*_1_(*y*_1_)^T^(*z*_0_ − *x*_1_) as ∥∇*F*_1_(*y*_1_) ∥_2_ ·∥*z*_0_ − *x*_1_∥_2_ · cos(*θ*) for some *θ* ∈ [0, *π/*2) and applying the *L*_2_ norm on both sides yields, after applying the triangle inequality on the right hand side,

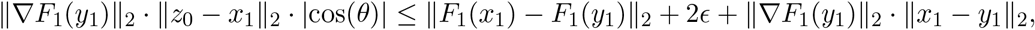

which after rearranging yields

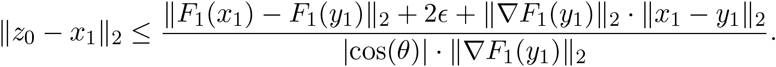

We write 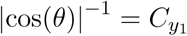 and note that *θ* is determined by *y*_1_ (and *z*) but independent of *x*_2_. Since *x*_1_ is fixed, and *F*_1_ is differentiable, it is also locally Lipschitz in a ball around *x*_1_ (note that the Lipschitz constant is independent of *x*_2_). Thus there exists 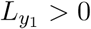 *>* 0 such that ∥ *F*_1_(*x*_1_) − *F*_1_(*y*_1_) ∥_2_ ≤ 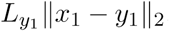. This is true for any *y*_1_ ∈ *S*_*δ*_(*x*_1_), thus by taking the supremum over *y*_1_ ∈ *S*_*δ*_(*x*_1_) we arrive at two constants *C*_*δ*_ *>* 0 and *L*_*δ*_ *>* 0 (independent of *y*_1_) which satisfy

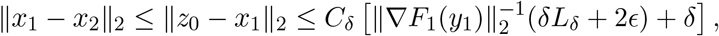

where it was used that ∥ *x*_1_ − *y*_1_ ∥_2_ = *δ* since *y*_1_ ∈ *S*_*δ*_(*x*_1_), and that ∥ *x*_1_ − *x*_2_∥_2_ ≤ ∥*x*_1_ − *z*_0_∥_2_.

The bound ∥ *x*_1_ − *x*_2_ ∥ _2_ ≤ ∥ *z*_0_ − *x*_1_ ∥ _2_ is conditional on *y*_1_ and *x*_1_, *x*_2_ being collinear. Since *x*_2_ is unknown, we take the supremum over all *y*_1_ ∈ *S*_*δ*_(*x*_1_), resulting in

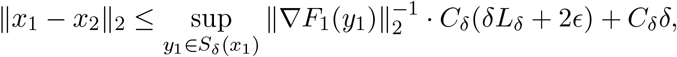

thus establishing the claimed upper bound. □

### C Additional results on simulated data

This section presents additional simulations using the simulation setup of Section 3.1.

#### C.1 Fused Lasso on simulated data

The fused Lasso (Tibshirani et al., 2005) is provided in the *fusedlasso* function of the R-package *genlasso* (Arnold & Tibshirani, 2020).

For the fused Lasso, a dependency structure among the entries of the parameter vector *β* ∈ ℝ^*p*^ has to be generated. This dependency structure is given by the adjacency matrix *E*, see Section 2.4. We generate *E* at random with edge probability 0.5 in each repetition. In this experiment, we employ the choices of *u, v*, and *w*_*j*_, *j* ∈ {1, *…, p*}, for the fused Lasso given in Section 2.4.

Figure 6 shows scaling results in *n* while keeping *p* = 100 fixed. We observe that the function fusedlasso of the R-package genlasso seems to have troubles finding good estimates, while optimizing the objective function of the fused Lasso, given in Section 2.4, with our progressive smoothing algorithm works much better and yields stable regression estimates having a low *L*_2_ norm and a high correlation with the truth. The *fusedlasso* function and minimizing the unsmoothed objective yield fastest results, followed by our smoothing approach. Progressive smoothing is a constant factor slower as expected. Overall, there only seems to be a weak runtime dependence on *n*.

**Figure 6:**
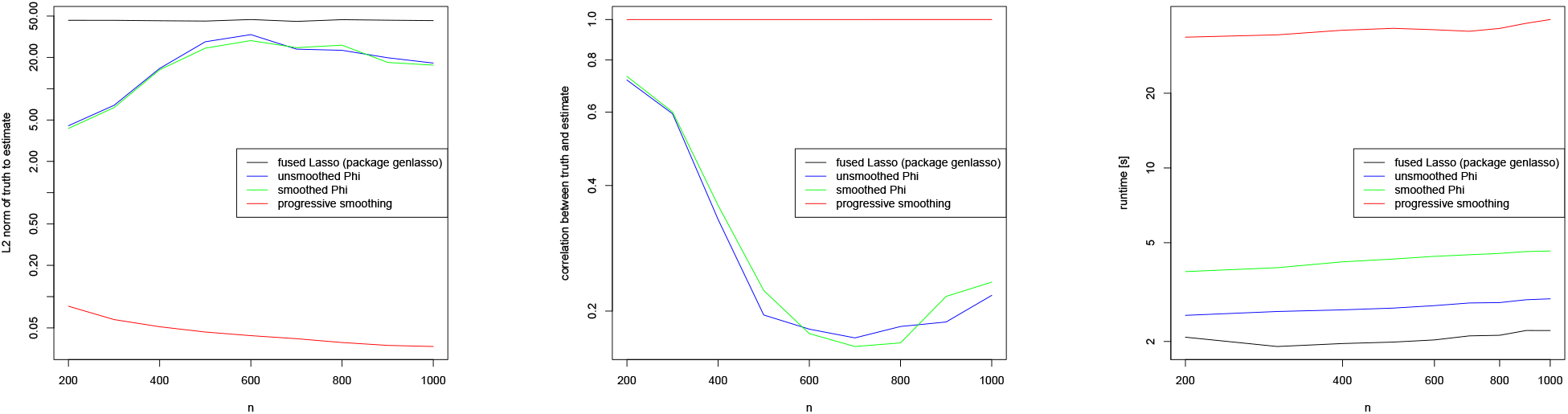
Fused Lasso: *L*_2_ norm (left) and correlation (middle) between simulated truth and parameter estimates as a function of *p* while *n* = 100. Runtime in seconds (right) has a log scale on both axes. The comparison includes fusedlasso (R-package genlasso), unsmoothed Φ, smoothed surrogate Φ^*μ*^, and progressive smoothing.

Figure 7 shows similar results for the scaling in *p*. Here, the *fusedlasso* function again yields estimates with the largest deviation from the generated truth in the *L*_2_ norm, and also the lowest correlation with the true parameters. Progressive smoothing yields the most accurate and stable results with respect to both *L*_2_ norm and correlation, while the other two approaches (unsmoothed objective and smooth surrogate, denoted as unsmoothed Phi and smoothed Phi) exhibit a more unstable behavior. The runtime scaling of all methods is roughly similar in *p*.

**Figure 7:**
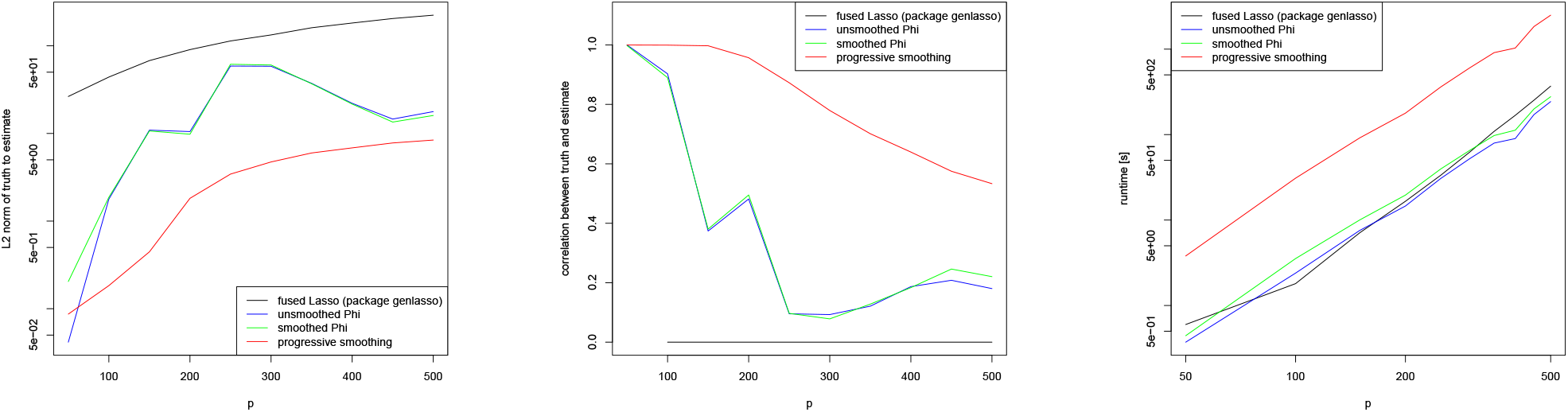
Fused Lasso: *L*_2_ norm (left) and correlation (middle) between simulated truth and parameter estimates as a function of *p* while *n* = 100. Runtime in seconds (right) has a log scale on both axes. The comparison includes fusedlasso (R-package genlasso), unsmoothed Φ, smoothed surrogate Φ^*μ*^, and progressive smoothing.

### C.2 Graphical Lasso on simulated data

The graphical Lasso (Friedman et al., 2008) is provided in the *glasso* function of the R-package *glasso* (Friedman et al., 2019).

For the graphical Lasso, we generate a sample covariance matrix *S* from a Wishart distribution with sample size 1, *p* ∈ {5, *…*, 30} degrees of freedom, and scale matrix set to the *p* × *p* dimensional identity matrix, see Section 2.4. To define Φ and its smooth surrogate Φ^*μ*^, we employ the functions *u, v*, and *w*_*j*_, *j* ∈ {1, *…, m*} with *m* = *p*(*p* + 1)*/*2, for the graphical Lasso given in Section 2.4.

We then compute estimates for the unsmoothed graphical Lasso objective function using the R-package glasso. Additionally, we minimize the unsmoothed Φ for the graphical Lasso as well as our smooth surrogate Φ^*μ*^ using the BFGS algorithm in R’s function *optim*, and employ our progressive smoothing algorithm. To optimize over all positive definite matrices Θ in the objective function of the graphical Lasso (see Section 2.4), we parametrize Θ with its Cholesky decomposition. To be precise, we write Θ = *CC*^T^, where *C* is a lower triangular matrix. This ensures that Θ will always be positive definite. As Θ is fully parametrized via *C*, and *C* is lower diagonal, we actually only optimize over *p*(*p* + 1)*/*2 parameters.

We report the correlation between the true and estimated precision matrices (for this the matrices are unfolded as vectors), as well as the runtime.

Results are given in Figure 8. The results show that estimates of the precision matrix found with the glasso R-package and by minimizing the unsmoothed objective function have a lower correlation with the generated truth than the estimates obtained with our two smoothing approaches. The implementation of the graphical Lasso in the R-package glasso seems highly optimized, resulting in fast runtimes. Our smoothing approaches are not as highly optimized and thus slower than glasso, though all three exhibit roughly a similar runtime scaling.

**Figure 8:**
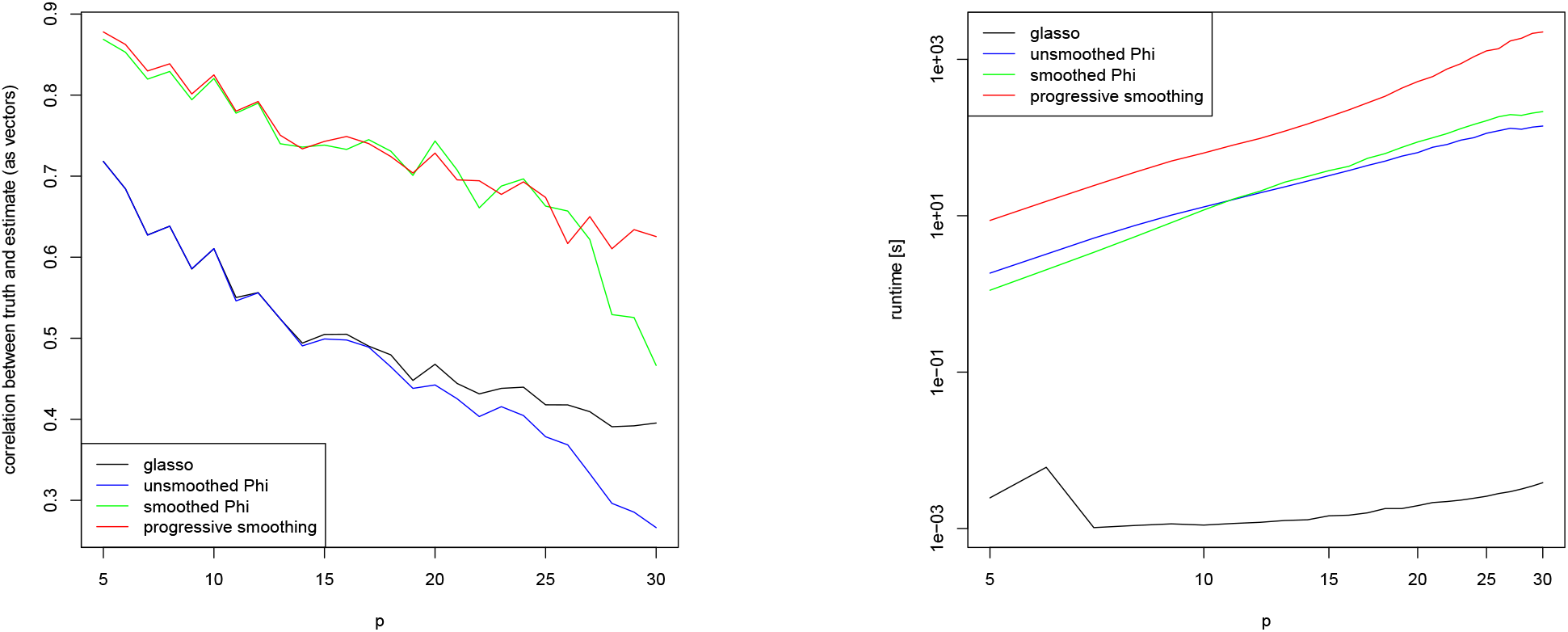
Graphical Lasso: correlation (left) between simulated truth and parameter estimates as a function of *n*. Runtime in seconds (right) has a log scale on both axes. The comparison includes glasso (R-package glasso), unsmoothed Φ, smoothed surrogate Φ^*μ*^, and progressive smoothing.

## Notes

### Competing Interest Statement

The authors have declared no competing interest.

### Summary of Updates

Simulations are updated.

## References

Arnold, T. B. & Tibshirani, R. J. (2020). genlasso: Path Algorithm for Generalized Lasso Problems. R-package version 1.5: https://cran.r-project.org/package=genlasso.

Banerjee, O., Ghaoui, L. E. & d’Aspremont, A. (2008). Model Selection Through Sparse Maximum Likelihood Estimation for Multivariate Gaussian or Binary Data. Journal of Machine Learning Research 9, 485–516.

Beck, A. & Teboulle, M. (2009). A Fast Iterative Shrinkage-Thresholding Algorithm for Linear Inverse Problems. SIAM J Imaging Sciences 2, 183–202.

Beck, A. & Teboulle, M. (2012). Smoothing And First Order Methods: A Unified Framework. Siam J Optim 22, 557–580.

Becker, S. R. & Candès, E. J. (2011). Templates for convex cone problems with applications to sparse signal recovery. Math Prog Comp 3, 165–218.

Belloni, A., Chernozhukov, V. & Wang, L. (2011). Square-root lasso: pivotal recovery of sparse signals via conic programming. Biometrika 98, 791–806.

Chen, X., Kim, S., Lin, Q., Carbonell, J. G. & Xing, E. P. (2010a). Graph-Structured Multi-task Regression and an Efficient Optimization Method for General Fused Lasso. 1005.3579 1–21.

Chen, X., Lin, Q., Kim, S., Carbonell, J. G. & Xing, E. P. (2010b). An efficient proximal gradient method for general structured sparse learning. Journal of Machine Learning Research 11.

Chen, X., Lin, Q., Kim, S., Carbonell, J. G. & Xing, E. P. (2012). Smoothing proximal gradient method for general structured sparse regression. Ann Appl Stat 6, 719–752.

Chi, E., Goldstein, T., Studer, C. & Baraniuk, R. (2018). fasta: Fast Adaptive Shrinkage/Thresholding Algorithm. R-package version 0.1.0: https://cran.r-project.org/package=fasta.

Daubechies, I., Defrise, M. & Mol, C. (2004). An iterative thresholding algorithm for linear inverse problems with a sparsity constraint. Comm. Pure Appl. Math. 57, 1413–1457.

Dondelinger, F. & Mukherjee, S. (2020). The joint lasso: high-dimensional regression for group structured data. Biostatistics 21, 219–235.

Efron, B., Hastie, T., Johnstone, I. & Tibshirani, R. (2004). Least angle regression. Ann Stat 32, 407–499.

Fan, J. & Li, R. (2001). Variable Selection via Nonconcave Penalized Likelihood and its Oracle Properties. J Am Stat Assoc 96, 1348–1360.

Friedman, J., Hastie, T. & Tibshirani, R. (2008). Sparse inverse covariance estimation with the graphical lasso. Biostatistics 9, 432–441.

Friedman, J., Hastie, T. & Tibshirani, R. (2010a). Regularization Paths for Generalized Linear Models via Coordinate Descent. Journal of Statistical Software 33, 1–22.

Friedman, J., Hastie, T. & Tibshirani, R. (2010b). Regularization Paths for Generalized Linear Modelsvia Coordinate Descent. J Stat Softw 33, 1–22.

Friedman, J., Hastie, T. & Tibshirani, R. (2019). glasso: Graphical Lasso: Estimation of Gaussian Graphical Models. R-package version 1.11: https://cran.r-project.org/package=glasso.

Friedman, J., Hastie, T., Tibshirani, R., Narasimhan, B., Tay, K., Simon, N. & Qian, J. (2020). glmnet: Lasso and Elastic-Net Regularized Generalized Linear Models. R-package version 4.0: https://cran.r-project.org/package=glmnet.

Hahn, G., Banerjee, M. & Sen, B. (2017). Parameter Estimation and Inference in a Contin-uous Piecewise Linear Regression Model. http://www.cantab.net/users/ghahn/preprints/PhaseRegMultiDim.pdf.

Hahn, G., Lutz, S., Laha, N., Cho, M., Silverman, E. & Lange, C. (2021). A fast and efficient smoothing approach to LASSO regression and an application in statistical genetics: polygenic risk scores for Chronic obstructive pulmonary disease (COPD). Stat Comput 31, 1–11.

Hahn, G., Lutz, S. M., Laha, N. & Lange, C. (2020). smoothedLasso: Smoothed LASSO Regres-sion via Nesterov Smoothing. R-package version 1.4: https://cran.r-project.org/package=smoothedLasso.

Haselimashhadi, H. & Vinciotti, V. (2016). A Differentiable Alternative to the Lasso Penalty. 1609.04985 1–12.

Hastie, T. & Efron, B. (2013). lars: Least Angle Regression, Lasso and Forward Stagewise. R-package version 1.2: https://cran.r-project.org/package=lars.

Massias, M., Fercoq, O., Gramfort, A. & Salmon, J. (2018). Generalized Concomitant Multi-Task Lasso for Sparse Multimodal Regression. In Proceedings of the 21st International Conference on Artificial Intelligence and Statistics (AISTATS) 2018, Lanzarote, Spain, vol. 84, PMLR.

Mazumder, R., Choudhury, A., Iyengar, G. & Sen, B. (2019). A Computational Framework for Multivariate Convex Regression and Its Variants. J Am Stat Assoc 114, 318–331.

Michelot, C. (1986). A finite algorithm for finding the projection of a point onto the canonical simplex of Rn. J Optimiz Theory App 50, 195–200.

Ndiaye, E., Fercoq, O., Gramfort, A., Leclère, V. & Salmon, J. (2017). Efficient Smoothed Con-comitant Lasso Estimation for High Dimensional Regression. In 7th International Conference on New Computational Methods for Inverse Problems.

Nesterov, Y. (1983). A method of solving a convex programming problem with convergence rate O(1/k^2^). Dokl Akad Nauk SSSR 269, 543–547.

Nesterov, Y. (2005). Smooth minimization of non-smooth functions. Math. Program. Ser. A 103, 127–152.

R Core Team (2014). R: A Language and Environment for Statistical Computing. R Foundation for Stat Comp, Vienna, Austria.

Seijo, E. & Sen, B. (2011). A continuous mapping theorem for the smallest argmax functional. Electron J Stat 5, 421–439.

Tibshirani, R. (1996). Regression Shrinkage and Selection Via the Lasso. J Roy Stat Soc B Met 58, 267–288.

Tibshirani, R., Saunders, M., Rosset, S., Zhu, J. & Knight, K. (2005). Sparsity and Smoothness via the Fused Lasso. J Roy Stat Soc B Met 67, 91–108.

Wang, W. & Carreira-Perpiñán, M. (2013). Projection onto the probability simplex: An efficient algorithm with a simple proof, and an application. 1309.1541 1–5.

Yuan, M. & Lin, Y. (2005). Model selection and estimation in regression with grouped variables. J Roy Stat Soc B Met 68, 49–67.

Zou, H. & Hastie, T. (2005). Regularization and variable selection via the elastic net. J Roy Stat Soc B Met 67, 301–320.

